# Sex-specific brain effective connectivity patterns associated with negative emotions

**DOI:** 10.1101/2024.04.02.587489

**Authors:** Tajwar Sultana, Dua Ijaz, Fareha Asif Khan, Maryam Misaal, Elvisha Dhamala, Adeel Razi

## Abstract

Sex differences in effective brain connectivity in emotional intelligence, emotional regulation, and stimuli-induced negative emotions have been highlighted in previous research. However, to our knowledge, no research has yet investigated the sex-specific effective connectivity related to negative emotions in healthy population during resting-state. The goal of this study is to find the association between sex-specific resting-state effective brain connectivity and basic negative emotions. For this, we have employed the NIH emotion battery of the three self-reported, basic negative emotions — anger-affect, fear-affect, and sadness which we divided into high, moderate, and low emotion scores in each. The dataset comprises 1079 subjects (584 females) from HCP Young Adults. We selected large-scale resting-state brain networks important for emotional processing namely default mode, executive, and salience networks. We employed subject-level analysis using spectral dynamic causal modelling and group-level association analyses using parametric empirical Bayes. We report association of the self-connection of left hippocampus in females in high anger-affect, fear-affect, and sadness, whereas in males we found involvement of dorsal anterior cingulate cortex (dACC) in all three negative emotions - association of right amygdala to dACC in high anger-affect, association of the self-connection of dACC in high fear-affect, and association of dACC to left hippocampus in high sadness. Our findings primarily revealed the effective brain connectivity that is related to the higher levels of negative emotions that may lead to psychiatric disorders if not regulated. Sex-specific therapies and interventions that target psychopathology can be more beneficial when informed by the sex-specific resting-state effective connectivity.

## Introduction

Does the male and female brain have different brain connectivity for emotions? This question is of utmost importance in recent times, as sex is a crucial aspect of human identity and has been the subject of numerous studies and research over the years. The National Institute of Mental Health, 2011 workshop on “Sex Differences in Brain, Behavior, Mental Health and Mental Disorders” highlighted the necessity of incorporating sex differences or sex as a variable in research and clinical practices (McCarthy et al., 2012). Studies have explored sex differences in emotions, and while the findings are not entirely consistent across all studies, some patterns have emerged. Research has shown that males and females tend to experience emotions differently, and there are documented sex differences in emotional expression (Brody, 1997; Burt, 2014; Deng et al., 2016; Ficek-Tani et al., 2023; Fischer & Evers, 2011; Gross & John, 2003). Males and females also exhibit diverse patterns of emotional responses to various stimuli (Bradley et al., 2001; Lithari et al., 2010).

Negative emotions are feelings that are generally unpleasant and associated with unpleasant experiences, such as sadness, anger, fear, disgust, and frustration (Deng et al., 2016). Negative emotions serve an important adaptive function, as they help us respond to and deal with challenging or threatening situations (de Marco et al., 2006). These emotions are felt by both men and women, regardless of sex yet females usually express stronger negative emotions as compared to males (Deng et al., 2016), which may explain why females are more prone to mood disorders (Brody, 1997; Burt, 2014; Deng et al., 2016; Fischer & Evers, 2011; Gross & John, 2003; Hess et al., 2000). In previous research, it has been suggested that differences in brain anatomy and function contribute to these variances between males and females (Cosgrove et al., 2007).

The brain pathways associated with emotion processing were previously explored through investigations of functional and effective connectivity. Stronger functional connectivity in a cingulo-opercular network was linked to better suppression in females during the regulation of negative emotion, whereas males showed stronger functional connectivity in posterior areas of the ventral attentional network (Stoica et al., 2021). In another study, sex differences in functional connectivity during the unattended processing of facial expressions were investigated. The results showed that females exhibited stronger and more widely distributed brain activation and connectivity compared to males (J. Zhang et al., 2018). Previous effective connectivity study having stimuli induced negative emotions concluded that men have a much higher granger causal connectivity from the right amygdala than from the dorsolateral medial prefrontal cortex which suggested that men have more evaluative brain responses than just affective when processing negative emotions (Lungu et al., 2015). Various other studies have also explored sex differences in response to emotion regulation (Domes et al., 2010; Koch et al., 2007; Mak et al., 2009; McRae et al., 2008; Stoica et al., 2021), emotion reactivity (Domes et al., 2010; Fine et al., 2009; McClure et al., 2004; McRae et al., 2008; Shirao et al., 2005; Thomas et al., 2001; L. M. Williams et al., 2005), emotion experience (Burt, 2014; Deng et al., 2016; Fischer & Evers, 2011; Gross & John, 2003; Hess et al., 2000) and emotion perception (Fischer et al., 2018; Han et al., 2008; Kempton et al., 2009; Killgore & Yurgelun-Todd, 2001; Lopez-Zafra & Gartzia, 2014; Wildgruber et al., 2002). Ignoring these distinctive outcomes between males and females may result in unsuccessful interventions and therapies for emotional illnesses such as anxiety and depression. Additionally, failing to acknowledge how men and women process emotions differently can reinforce negative sex stereotypes (Fischer, 1993; Heesacker et al., 1999; Lopez-Zafra & Gartzia, 2014) and contribute to the ongoing sex disparities in access to mental health care. For instance, sex stereotypes may encourage the notion that men shouldn’t be vulnerable or emotional, which could result in a lack of recognition and treatment of male emotional problems (Fischer et al., 2018; Lopez-Zafra & Gartzia, 2014; McKenzie et al., 2022).

Previous research has been focused on examining brain functional or effective connectivity while emotions were induced using stimulus in the scanner. This induction mechanism may reflect transient emotions or state emotions but might not be a good indicator of trait-like emotions. The self-report measure of emotions usually assesses emotions based on the history of either a week or a fortnight or lifetime. Questionnaires based on lifetime experiences assess personality traits based on emotions such as neuroticism while those based on narrower past window may be used to assess trait-like personality. There is a lack of sex-specific brain connectivity research examining trait-like personality based on negative emotions. Therefore, current work is based on the self-report scores of negative emotions in which the subjects reported their feelings during the past week.

Although there has been extensive research on sex differences in emotions, neuroscientific exploration of the sex differences in resting-state effective brain connectivity specifically related to basic negative emotion is much required. To investigate this gap comprehensively, we expanded upon the previous studies and investigated the underlying neural mechanisms of emotional trait-like personality in males and females using resting-state fMRI. Resting-state fMRI allows for the examination of intrinsic brain activity and connectivity patterns without the need for any external stimuli. Based on the established literature, we anticipate significant associations of the amygdala connectivity in anger in males along with associations with the connectivity of insula, hippocampus, and medial prefrontal cortex (mPFC), while in females, we expect strong associations with the connectivity of hippocampus, dorsal anterior cingulate cortex (dACC), amygdala, and mPFC in fear and sadness. Previous literature findings suggests that males are often perceived as more susceptible to anger, while females are more prone to fear and sadness (Fahlgren et al., 2022; Fischer & Evers, 2011; Sadeh et al., 2011; Sharkin, 1993). Anger has been consistently linked with the amygdala (Birnbaum et al., 1980; Carlson et al., 2010; Swartz et al., 2019), whereas fear and sadness have been associated with the amygdala, mPFC, and dACC (Blair, 2012; Jamieson et al., 2021). Another study found that women have significantly higher levels of activity in those brain regions which are responsible for continuously evaluating emotional experience and subsequently remembering the stimuli that elicited the strongest emotions, hence, demonstrating that sex differences also exist in emotional memory (Canli et al., 2002). The hippocampus’s function in the emotion-memory processing is a matter of well-established opinion (Bisby & Burgess, 2017; Treves & Rolls, 1994). Therefore, it is reasonable to expect sex-specific associations of effective brain connectivity involving hippocampus with basic negative emotions.

## Methods

### fMRI Dataset

The dataset used in our research was acquired from Connectome Coordination Facility (CCF) (https://www.humanconnectome.org/). CCF is an online platform that provides data from several studies of the human brain, which are publicly available. These studies are known as Human Connectome Projects (HCP) (Elam et al., 2021). Our dataset is from healthy adult connectomes that is HCP Young Adult which was preprocessed using fMRI volume preprocessing pipeline (Glasser et al., 2013). We considered 1079 subjects (Male = 495, female = 584; age = 22-35) for analysis. The resting-state fMRI data was acquired using 3T MRI scanner. The acquisition parameters for resting state images were repetition time (TR) = 720 ms, Echo time (TE) = 3.31 ms, flip angle = 54°, FOV = 208 ˣ 180 mm. The resting-state fMRI data was collected in four runs of 15 minutes each, two in one session and two in the next session, with 1200 frames per run and 72 slices covering the whole brain. We utilized only the functional data from the first run of each subject.

### Subjects

The large-scale dataset contains 1079 subjects from the HCP. This participant pool was divided into two groups: males and females, with high, moderate, and low emotion scores in each. While the scores of most of the subjects followed a normal distribution pattern, there was a discrepancy: some subjects displayed the same score within each emotion group. The bimodal distribution that resulted from these minimal self-reported scores was particularly noticeable in the low anger-affect, fear-affect, and sadness. The use of approximations during score calculations might be the cause of this discrepancy. In accordance with the technique laid out by (Bouziane et al., 2022), in order to rule out any possible anomalies, we removed all the subjects showing the same score. This adjustment resulted in total of 176 subjects (85 male subjects and 91 female subjects) being removed from the original counts. This left us with 410 male subjects and 493 female subjects. Please see Figure S1 for the distribution of subjects across the sex and various emotion categories.

### Emotional assessment

The emotional assessment of HCP subjects was done using NIH toolbox emotion battery (https://www.nia.nih.gov/research/resource/nih-toolbox). The subjects emotional scores were collected through self-report using Computer Adaptive Test (CAT). A 5-point scale with options ranging from “never” to “always” was used for each administered item. Item Response Theory (IRT) techniques were used to grade the survey. For each participant, an IRT-derived theta score was produced which was transformed into an uncorrected standard score (T-Score). T-scores that are T ≥ 60 indicate strong levels of emotions, while T-scores that are T ≤ 40 indicate low levels of emotions. We used three self-reported basic negative emotions that are anger-affect (evaluates the feeling of anger), fear-affect (evaluates anxiety and fear) and sadness (evaluates unfavorable mood, negative self-perceptions) from the NIH Toolbox emotion battery.

### Regions of interest Selection

The regions of interest (ROIs) were taken from three large-scale resting state brain networks that are well known and have a high significance in emotional processing, namely: default mode network (DMN), executive network (EN), and salience network (SN). The cardinal brain regions that were chosen for DMN are: posterior cingulate cortex (PCC), medial prefrontal cortex (mPFC) and bilateral hippocampus (lHP and rHP), those selected from SN are: dorsal anterior cingulate cortex (dACC), bilateral anterior insula (lAI and rAI) and bilateral amygdala (lAMG and rAMG) and those that belong to EN are: bilateral dorsal lateral prefrontal cortex (lDLPFC and rDLPFC). These regions of interest were acquired from our previous study that investigated the underlying neural mechanism of neuroticism and self-reported basic negative emotions (Sultana et al., 2023).

### Data analysis

The data analysis was done using SPM12 (Welcome Trust Centre for Neuroimaging, London, UK; code available at: https://github.com/spm/spm12). In first level analysis spectral DCM (spDCM: (Friston et al., 2014; Novelli et al., 2024; Razi et al., 2015; Razi & Friston, 2016)) was used to determine the resting-state effective connectivity between different brain regions. A fully connected DCM model (having both intrinsic and extrinsic connections) over 11 ROIs were estimated for each subject. Then in the second level analysis, associations were found between estimated model and self-reported scores in which estimated effective connectivity for each subject was taken to group-level to estimate group effects. The HCP subjects were divided into two groups (males and females) which were subdivided according to their score levels (high, moderate, and low) for each emotion category (anger-affect, fear-affect, and sadness) that makes a total of 18 set of analysis. The group-level associations with emotion scores were determined using parametric empirical Bayes (PEB) framework (Friston et al., 2016; Zeidman et al., 2019) that specify hierarchical model of connectivity parameters with group effects as covariates. It is a Bayesian GLM with a design matrix defined with regressors - first column of all ones that define the commonalities in connectivity across all subjects and the second column of mean-centered emotion scores for the association analysis. Bayesian model reduction (BMR) was then used to score reduced models, nested within the fully connected mode, that were examined using free energy as a proxy for the model evidence. Bayesian model averaging (BMA) was then performed to average the parameters of models weighted by their model evidence. This resulted in group-level parameters that determined the association between effective connectivity and emotion scores of the group. The whole methodology is summarized in Figure 1.

**Figure 1.**
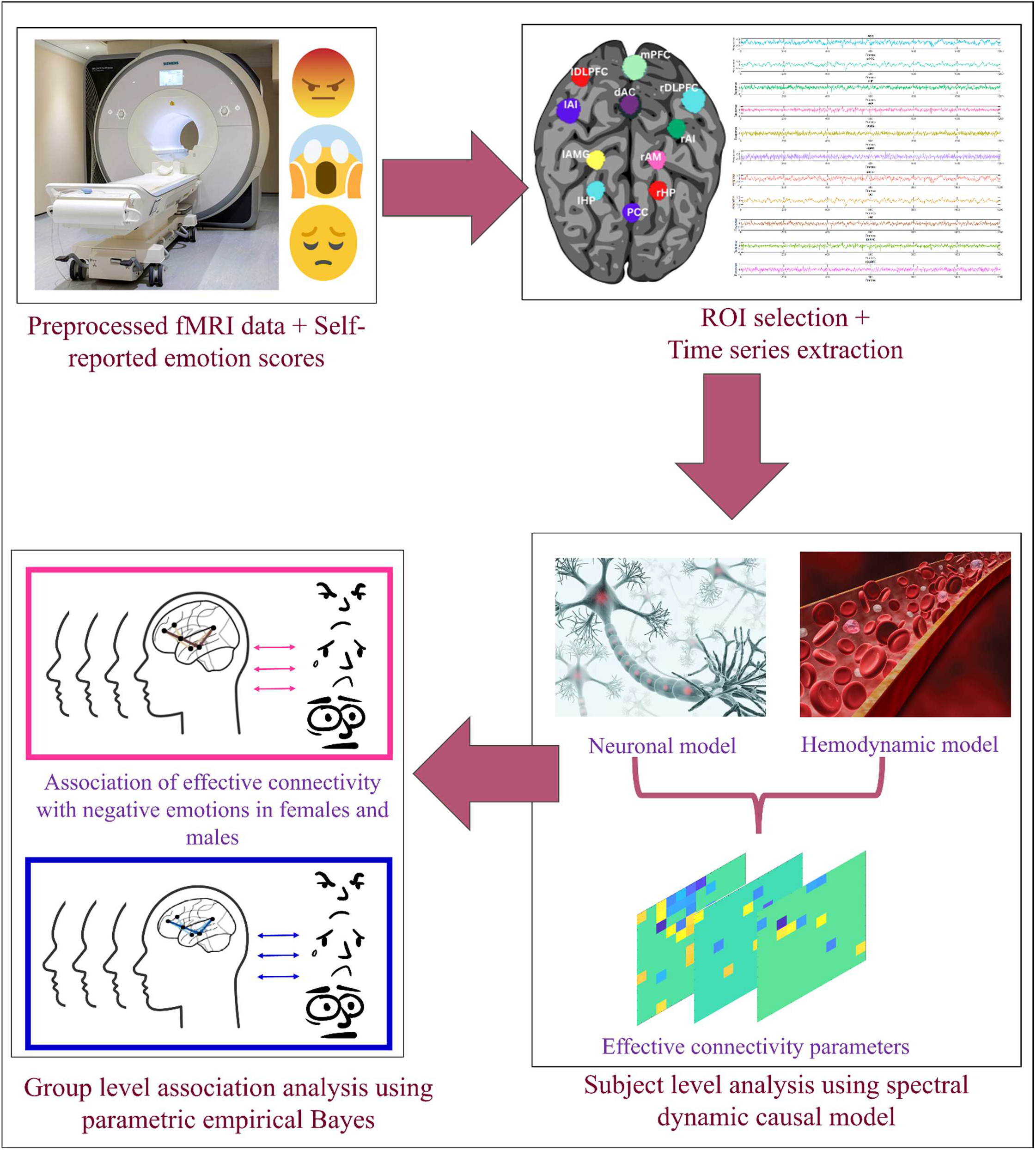
**Methodology flow chart** This flowchart illustrates the methodology. The process starts with preprocessed fMRI dataset and self-reported basic negative emotion scores followed by ROI selection and time series extraction. Then subject-level analysis was conducted using spectral dynamic causal model followed by parametric empirical Bayes for group-level association analysis.

## Results

### Self-reported basic negative emotions

The self-reported scores of negative emotions: anger-affect, fear-affect, and sadness were divided into high, moderate, and low level for both males (M) and females (F) based on their T-scores, that is ≤ 40 for low level, 40 ≤ T ≤ 60 for moderate level and ≥ 60 for high level of emotion scores. Figure S2 reports the distributions with mean and standard deviation of each score for both males and females. The mean scores for each category are: low anger-affect (M = 36.68 ± 2.35, F = 37.11 ± 2.41), moderate anger-affect (M = 49.66 ± 4.74, F = 49.34 ± 5.04), high anger-affect (M = 65.03 ± 4.38, F = 64.11 ± 5.03), low fear-affect (M = 38.35 ± 1.43, F = 37.46 ± 1.01), moderate fear-affect (M = 50.23 ± 4.73, F = 51.26 ± 4.51), high fear-affect (M = 65.26 ± 4.04, F = 64.64 ± 4.71), low sadness (M = 38.91 ± 0.42, F = 39.16 ± 0.42), moderate sadness (M = 49.10 ± 4.59, F = 48.29 ± 4.57) and high sadness (M = 64.31 ± 4.00, F = 65.09 ± 5.59).

### First level analysis: Accuracy of DCM estimation

The cross spectral density (CSD) for each subject’s time series data was first calculated and then used as a data feature for fitting spDCM. The quality of the DCM model fitting is evaluated by calculating the variance explained for each subject. The range of variance explained was 50% to 97% with an average of 87.06 ± 4.28%. For male group, the range of explained variance was 68% to 97% with an average of 86.90 ± 4.34% whereas for female group, the range of explained variance was 50% to 97% with an average of 87.23 ± 4.22%. The distribution of DCM estimation accuracy is shown in Figure S3.

### Second level analysis: Effective connectivity associations with negative emotion scores

In group level analysis we analyzed the associations between effective connectivity and emotion scores using PEB. The results with statistically strong evidence i.e., posterior probability > 0.95 are reported only. The mean connectivity matrices, at the group level, that show inhibitory and excitatory connectivity are shown in Figures S4-S6. The association results for each negative emotion are given below:

#### High anger-affect

The number of connections associated with high anger-affect were comparatively higher in females than males. There was total 32 connections in females while only 11 connections in males. In females, 13 positive and 19 negative associations were found which include the strongest negative association of the self-connection of lHP and the strongest positive association of the self-connection of rHP.

In males, 10 positive associations and 1 negative association were found which includes strongest positive association of the self-connection of lDLPFC and strongest negative association of rAMG to dACC inhibitory connection. Please refer to Figure 2 for detailed associations between effective connectivity and emotion scores.

**Figure 2.**
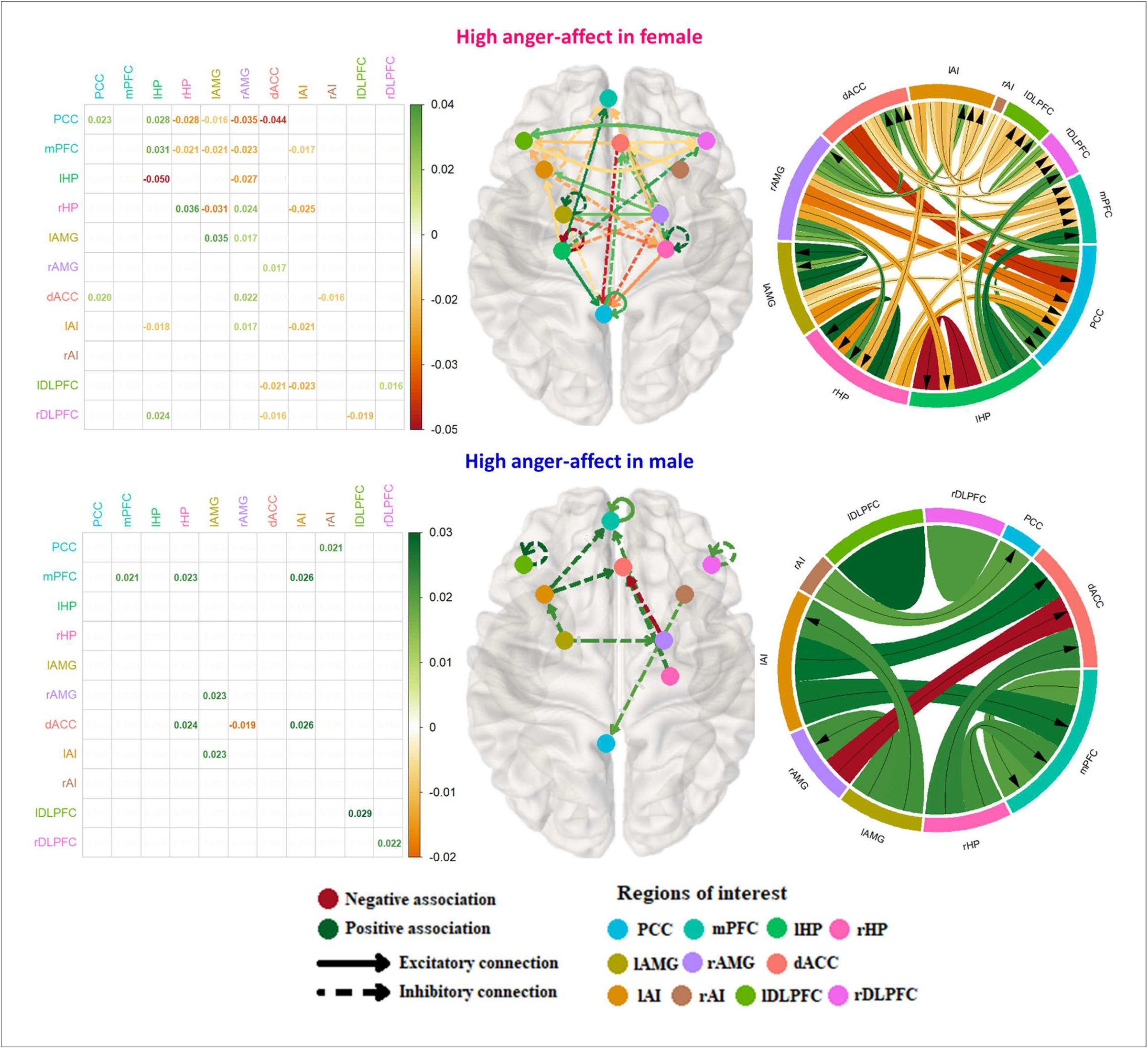
**Effective connectivity associations with high anger-affect scores** This figure illustrates the association between effective connectivity and high anger-affect scores in females (upper panel) and males (lower panel). In both figures the green shades represent the positive association and the red shade represents the negative association. The brain plot illustrates the connections between the regions where arrowhead refers to the region that is been influenced. The solid lines show the excitatory connection whereas the dotted lines illustrate inhibitory connections. A matrix on the bottom left side shows the standardized beta values of associations. The results are thresholded with posterior probability > 0.95. Abbreviations: mPFC = medial prefrontal cortex, PCC = posterior cingulate cortex, rHP = right hippocampus, lHP = left hippocampus, lAMG = left amygdala, rAMG = right amygdala, dACC = dorsal anterior cingulate cortex, lAI = left anterior insula, rAI = right anterior insula, lDLPFC = left dorsolateral prefrontal cortex, rDLPFC = right dorsolateral prefrontal cortex.

#### Low anger-affect

There was a total of 21 connections in females while 20 connections in males. In females, 10 positive and 11 negative associations were found which include the strongest negative association of the self-connection of lDLPFC and the strongest positive association of dACC to lDLPFC inhibitory connection.

In males, 10 positive and 10 negative associations were found which include the strongest negative association of the self-connection of rAMG and the strongest positive association of the self-connection of lAMG. Please refer to Figure S7 for detailed associations between effective connectivity and emotion scores.

#### Moderate anger-affect

There was a total of 4 connections in females while 8 connections in males. In females, 4 positive and no negative associations were found which include the strongest positive association of the self-connection of rHP.

In males, 2 positive and 6 negative associations were found which include the strongest negative association of the self-connection of mPFC and the strongest positive association of the self-connection of rHP. Please refer to Figure S8 for detailed associations between effective connectivity and emotion scores.

#### High fear-affect

The number of connections associated with high fear-affect were comparatively higher in males than females. There was total 23 connections in males while only 10 connections in females. In females, 4 positive and 6 negative associations were found which include the strongest negative association of the self-connection of lHP and the strongest positive association of dACC to lAMG inhibitory connection.

In males, 10 positive and 13 negative associations were found which include the strongest positive association of the self-connection of dACC and the strongest negative association of the self-connection of lDLPFC. Please refer to Figure 3 for detailed associations between effective connectivity and emotion scores.

**Figure 3.**
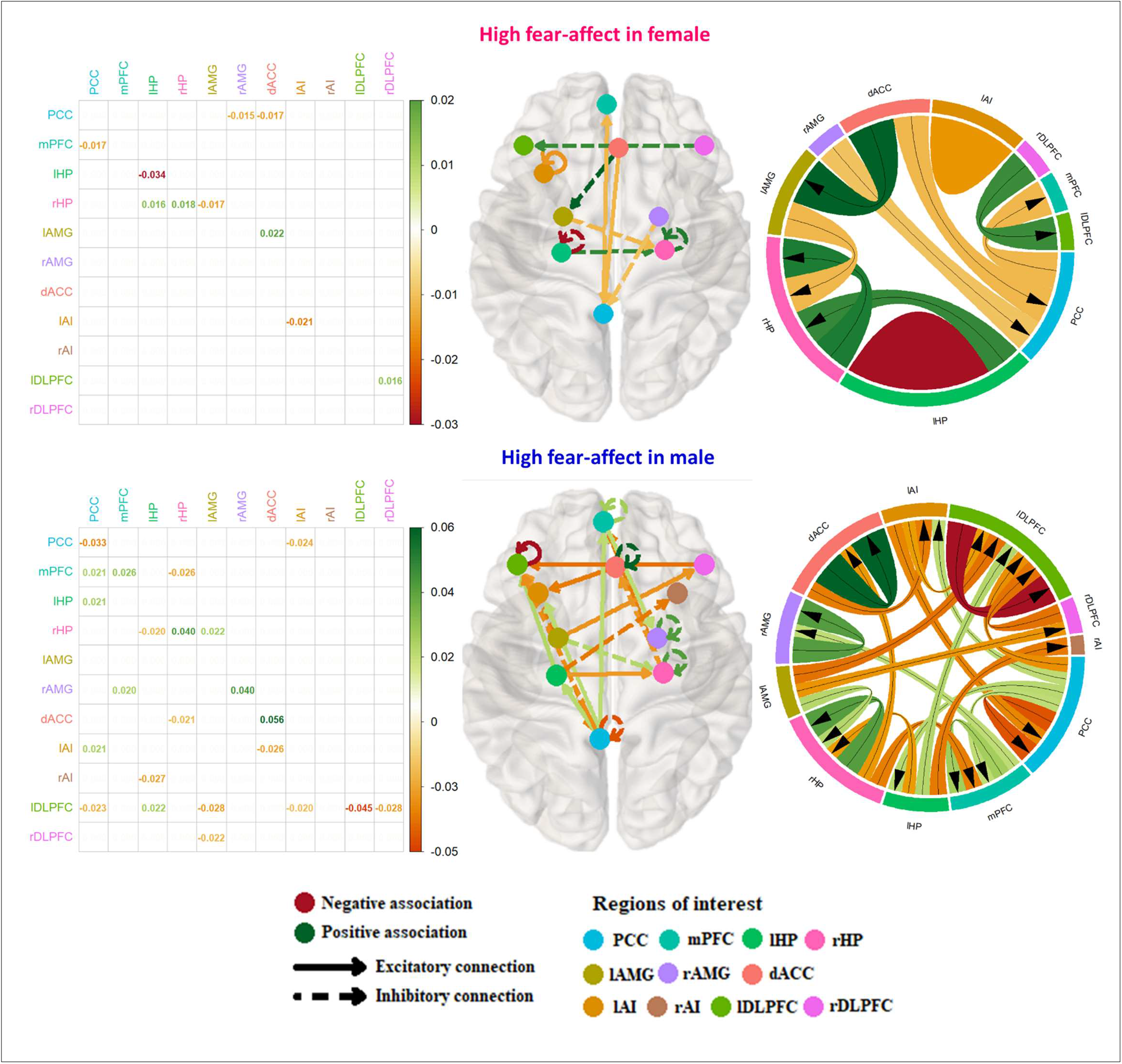
**Effective connectivity associations with high fear-affect scores** This figure illustrates the association between effective connectivity and high fear-affect scores in females (upper panel) and males (lower panel). The abbreviations and color arrangement are same as in Figure 2.

#### Low fear affect

There was a total of 26 connections in males and 22 connections in females. In females, 8 positive and 14 negative associations were found which include the strongest positive association of the self-connection of rHP and the strongest negative association of rAMG to PCC inhibitory connection.

In males, 13 positive and 13 negative associations were found which include the strongest positive association of the self-connection of dACC and the strongest negative association of rAMG to rHP inhibitory connection. Please refer to Figure S9 for detailed associations between effective connectivity and emotion scores.

#### Moderate fear affect

There was a total of 26 connections in males and 22 connections in females. In females, there was no negative association while 3 positive associations were found which include the strongest positive association of the self-connection of lAI.

In males, 2 positive and 3 negative associations were found which include the strongest positive association of the self-connection of rHP and the strongest negative association of the self-connection of lHP. Please refer to Figure S10 for detailed associations between effective connectivity and emotions scores.

#### High sadness

There was a total of 19 connections in females while 17 connections in males. In females, 7 positive and 12 negative associations were found which include the strongest negative association of the self-connection of lHP and the strongest positive association of the self-connection of dACC.

In males, 11 positive and 6 negative associations were found which include the strongest negative association of lHP to rHP excitatory connection and the strongest positive association of dACC to lHP inhibitory connection. Please refer to Figure 4 for detailed associations between effective connectivity and emotion scores.

**Figure 4.**
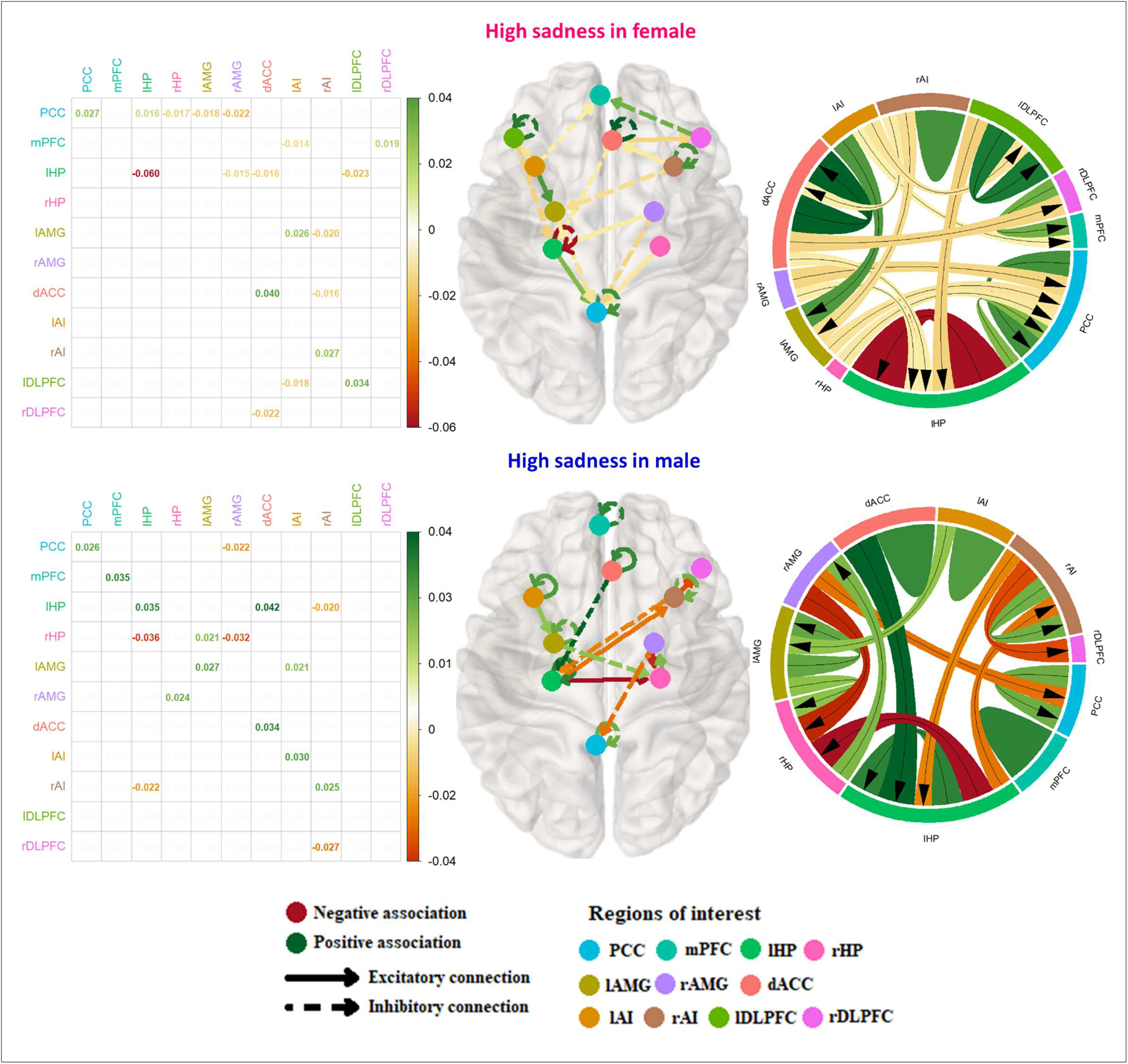
**Effective connectivity associations with high sadness scores** This figure illustrates the association between effective connectivity and high sadness scores in females (upper panel) and males (lower panel). The abbreviations and color arrangement are same as in Figure 2.

#### Low sadness

There was a total of 17 connections in males while 21 connections in females. In females, 15 positive and 6 negative associations were found which include the strongest positive association of the self-connection of lAMG and the strongest negative association of lAI to rAMG inhibitory connection.

In males, 11 positive and 6 negative associations were found which include the strongest positive association of rDLPFC to dACC excitatory connection and the strongest negative association of lHP to rAMG excitatory connection. Please refer to Figure S11 for detailed associations between effective connectivity and emotion scores.

#### Moderate sadness

There was a total of 8 connections in males while 4 connections in females. In females, there was no negative association while 4 positive associations were found which include the strongest positive association of the self-connection of lAI.

In males, 5 positive and 3 negative associations were found which include the strongest negative association of the self-connection of lHP and the strongest positive association of the self-connection of lDLPFC. Please refer to Figure S12 for detailed associations between effective connectivity and emotion scores.

## Discussion

Sex has a significant impact on basic negative emotions. Understanding the underlying causes for sex-specific self-reported basic negative emotions such as anger, fear and sadness is crucial to comprehend the attitudes of males and females towards negative emotions and their trait-like personality related to emotions. The goal of this study is to gain insight into sex-specific neural mechanisms in basic negative emotions over 11 regions of interest that are mostly involved in emotional processing, namely PCC, mPFC, dACC, bilateral AI, bilateral HP, bilateral AMG and bilateral DLPFC.

After conducting a thorough investigation that encompassed emotion scores in high, low, and moderate ranges, we choose to exclusively discuss the associations of high emotion scores. The decision has been driven by the notion that mental health illnesses are more likely to be exacerbated by high negative emotions (de Bles et al., 2019; Mahmud et al., 2021; Mouchet-Mages & Baylé, 2008). preceding research has also targeted the relationship of high negative emotions scores with mental disorders (Ng et al., 2022; Sahu et al., 2014; Yimam et al., 2022). Due to their greater significance, our discussion gives emphasis to the associations of high negative emotion scores.

The most highlighted and significant results (cf. Figure 5) are the strongest negative association of the self-connection of left hippocampus with high anger-affect, fear-affect and sadness in females and connectivity of dACC in all three negative emotions in males i.e., the strongest negative association of rAMG to dACC inhibitory connection in high anger-affect and the strongest positive association of the self-connection of dACC in high fear-affect and dACC to rHP inhibitory connectivity in high sadness. These findings provide support for our initial hypothesis, indicating that anger is indeed linked to amygdala connectivity in males. Furthermore, our results suggest that fear and sadness in females are strongly associated with the self-connection of the hippocampus, aligning with our initial predictions.

**Figure 5.**
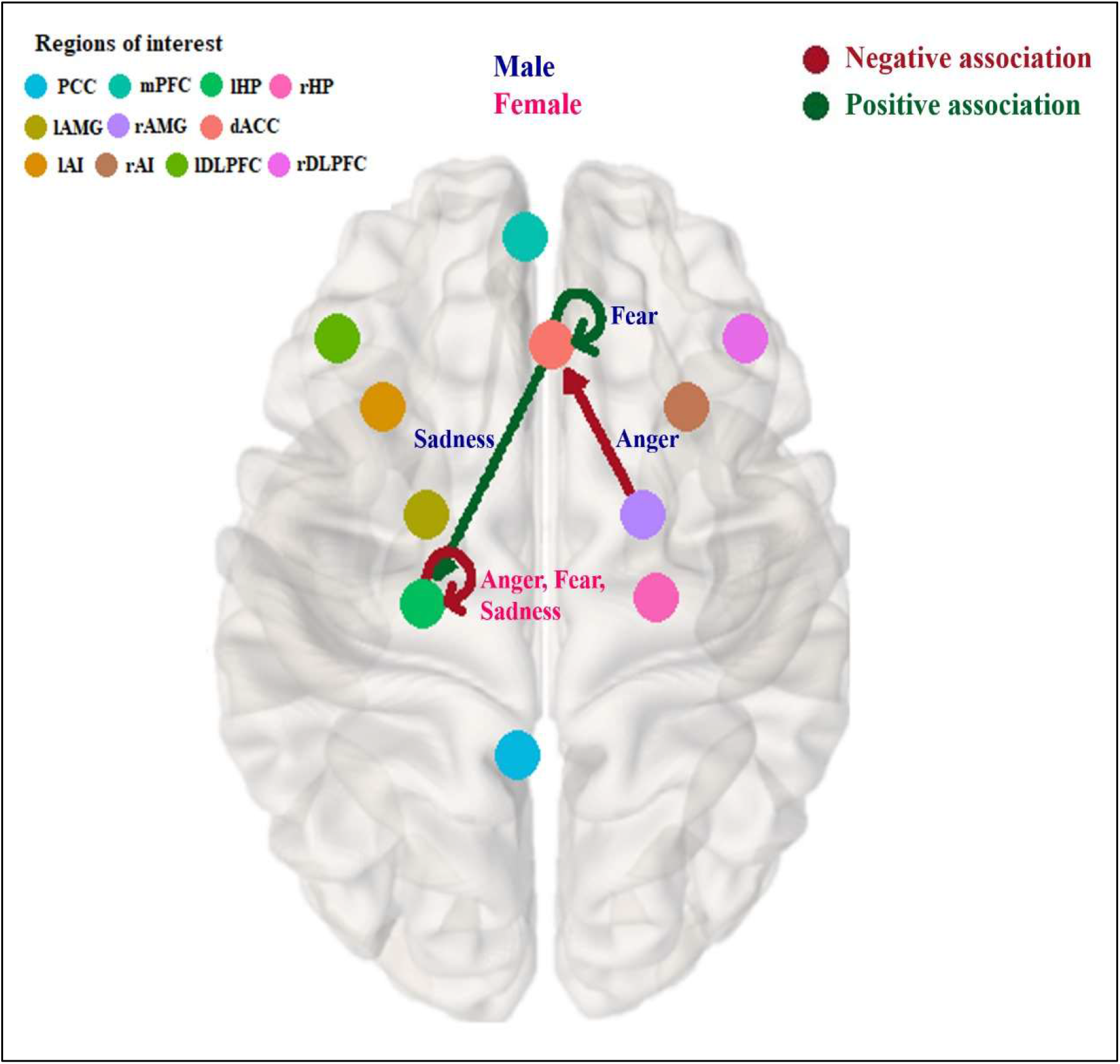
**The most highlighted outcomes of self-reported basic negative emotions** The red and green arrow illustrates positive and negative association respectively. The abbreviations and color arrangement are same as in Figure 2.

With strongest negative associations in all three negative emotions, the hippocampus stood out as a key finding in our study in females. Our results have shown the strongest negative association of the self-connection of hippocampus with each of the high-level negative emotions in females. Many previous studies have highlighted hippocampus as the key region found in emotion processing (Immordino-Yang & Singh, 2013; Zhu et al., 2019). Hippocampus is closely associated with memory, estrogen, and hippocampal volume (Taylor et al., 2020). Hippocampus and memory have a well-established role according to previous studies. A study for determining the role of hippocampus in memory suggested that hippocampus is closely involved in the formation, consolidation, and retrieval of episodic memories (Treves & Rolls, 1994) which is a type of long-term memory that involves the ability to recall specific events and experiences. Previous research has shown that hippocampal down regulation has been related with negative associative memory (Bisby et al., 2016). Hippocampal memory in females have also been highlighted in various previous studies. A previous study investigating sex differences in response to fear signals highlighted that female showed greater activation in hippocampus in response to fear stimuli (L. M. Williams et al., 2005). Another study revealed that hippocampus showed correlation with emotional arousal ratings and in the concurrence of emotional memory and emotional experience in females (Canli et al., 2002). This suggests that women’s hippocampal involvement in response to emotion is greater than that of men’s. and this may be the reason of the highlighted association of hippocampus with all three self-reported negative emotions in females. There have been many rodent-based studies trying to validate the more rigorous role of hippocampus in memory consolidation in females than males (Qi et al., 2016; Soler et al., 2021; Torromino et al., 2022; C. L. Williams & Meck, 1991). Hippocampal memory and female estrogen control have an established relation. In particular, estrogens are important hormones that regulate emotion, memory, mood, cognition and mental states, having a significant impact on physiology and behaviour (Fink et al., 1998; Luine, 2008). A prior study suggested that estrogen regulates and enhances hippocampal memory consolidation in females (Frick et al., 2018). According to previous research, women who underwent hormone therapy during menopause reported significantly greater hippocampal volumes compared to their peers who did not receive hormone therapy (Wnuk et al., 2012). These studies highlight the significance of estrogen in influencing the structure and function of the hippocampus in females. Some research studies have found sex differences in hippocampal volume (Nordenskjöld et al., 2015; Pintzka et al., 2015; Van Eijk et al., 2020). There are inconsistencies between studies, some reported larger hippocampus in males (Krogsrud et al., 2014; Lotze et al., 2019; Raz et al., 2004; Tamnes et al., 2018) others suggests that females have a larger hippocampus as compared to males (Kurth et al., 2017; Malykhin et al., 2017; Mueller et al., 2007), after accounting for individual differences. One study suggests that gray matter volume in the anterior-inferior hippocampus was larger in men than in women (Lotze et al., 2019). Although researchers have found significantly larger hippocampal volume in males than females but they recommend adjustment for total brain volume as no consistently evident sex difference or sex dimorphism exists in hippocampus size after adjusting for total brain volume (Perlaki et al., 2014; Ritchie et al., 2018; Tan et al., 2016).

Larger hippocampus volumes in older adults are associated with more rapid and accurate facial emotion recognition. Their preliminary findings imply that increased volume in the brain areas responsible for processing faces and emotions helps older people identify facial expressions of emotion more accurately (Szymkowicz et al., 2016). According to one study, males are found to have reduced functional connectivity in brain networks which involve the hippocampus as compared to females (Scheinost et al., 2015). In another study, 24 women with a history of recurrent major depression and 24 matched controls were assessed, MRI scans showed that the depressed women had smaller volumes of the hippocampus. The study suggested that the repeated stress experienced during depressive episodes might be the cause to cumulative damage to the hippocampus (Sheline et al., 1999). An overlap between the influence and the impact of emotions and memories is produced when amygdala and hippocampus interact or amygdala-hippocampus interaction creates overlap in emotions and memories (Phelps, 2004). These findings highlight the role of hippocampus in the complex interplay between sex, emotion, and memory.

Our findings’ significant hippocampal involvement may have been caused by the fact that some memories are linked to specific experiences and emotions. Therefore, we can infer that the participants’ memory must have been prompted by questions regarding the prior seven days and therefore this can be a reason for involvement of hippocampus connectivity in our results in both males and females.

According to our results, the effective connectivity of dACC is the most highlighted in males. For the regulation and processing of emotions, the role of anterior cingulate cortex is essential (Etkin et al., 2011; M. Zhang & Peng, 2023), and any disturbance in it can have a substantial effect. In another study, dACC was associated with social rejection and social distress (Eisenberger et al., 2003). In our study, the strongest negative association of the inhibitory connection from right amygdala to dACC was found with high levels of anger-affect. The amygdala is well known for its role in the processing of negative emotions, particularly anger (Blair, 2012; Carlson et al., 2010; Swartz et al., 2019). Another study found that the amygdala is involved in reactive aggression, caused by a perceived threat or harm (Bertsch et al., 2020). Deficits in impulse control are associated with behavioral problems like aggression. The term “impulsivity” refers to a tendency to act spontaneously and without consideration. It is more common in males than in females (Aron et al., 2007; Campbell & Muncer, 2009; Labouvie & McGee, 1986). One study suggest that males have an increased level of ACC activation than females (Liu et al., 2012) because male’s personality traits, like limited self-control, high impulsivity, and a tendency for negative emotionality raise the chance of impulsive aggression (Carver, 2005; Strüber et al., 2008). On the other hand, females were less impulsive and more likely to engage in defusing actions, making an effort to decrease the intensity of the anger-provoking, aggressive emotion (Beck & Bozman, 1995; Campbell & Muncer, 2008). This suggests inhibitory connection between right amygdala and dACC should be strengthened for better control of anger because decreasing inhibition would enhance anger. In light of these studies, our results might suggest that people with high anger-affect experience weakened connectivity from right amygdala to dACC, which could lead to difficulty controlling and managing these emotions. Poor causal connectivity from the right amygdala to dACC may lead to exaggerated or uncontrolled anger.

Our study also revealed a strong positive association between the self-connection of the dACC and high fear-affect in males. The dACC is well-known for its involvement in fear-related emotions (Milad et al., 2007; Sehlmeyer et al., 2011; Wang et al., 2023). It has previously been proposed that the dACC plays a role in the manifestation of fear in humans, potentially paving the way for the development of new treatments for anxiety in the future (Milad et al., 2007). Surprisingly, another study during the fear conditioning phase revealed that women exhibited significantly greater blood oxygen level-dependent (BOLD) signal changes in the right amygdala and ACC compared to men (Lebron-Milad et al., 2012). Another study explored how people who have high levels of trait anxiety have more activity in the amygdala and decreased activity in the dACC. Their findings imply that anxious individuals find it more difficult to overcome fear and may be more prone to anxiety disorders (Sehlmeyer et al., 2011).

In our study, the individuals experiencing high levels of sadness have a strong positive association of inhibitory connectivity from dACC to left hippocampus (lHP). This finding adheres with previous research that found ACC activation in response to sadness in healthy male participants (Koch et al., 2007). Furthermore, clinical research on the assessment and detection of sadness has emphasized sadness’s significant role in the development of depression (Mouchet-Mages & Baylé, 2008). It implies that if extreme sadness is not treated, it has the potential to progress into major depression. Notably, a study involving depressed males found stronger functional connectivity between the dACC and the left precuneus (Lee et al., 2018). This adds to the evidence that the dACC is involved in males with depressive disorders that can be due to high level of sadness.

## Conclusion

Our research provide evidence that the attitude towards basic negative emotions has different underlying neural mechanism in males and females. In males and females, the effective connectivity of dACC and left hippocampus play the most prominent role in heightened basic negative emotions respectively. Sex-specific therapies and interventions that target psychopathology can be more beneficial if they are informed by the resting-state effective connectivity associations of males and females.

## Future directions & limitations

In this study, only three basic negative emotions were investigated. Additional scores that are available in the HCP database such as aggression may be used in future analyses. Additionally, the participants are only young healthy adults of a particular age group. This study can be extended to involve participants from different age groups, hence inferring the sex-specific dynamics of brain connectivity in negative emotions across age-groups. The study is limited to the regions of interests spanning three large-scale resting-state networks, the sex-specific investigation of other resting-state networks will yield more insights into the brain dynamics of negative emotions.

