## Supplementary Information for "Sex-specific brain effective connectivity patterns associated with negative emotions"

### Dynamic causal modelling

A Bayesian framework called dynamic causal modeling (DCM) models effective connectivity using hemodynamic time series, which are blood oxygen level dependent (BOLD) data. The fMRI cannot directly measure the neural activity – it measures the changes in hemodynamics response and then infer the neural activity. A two-component generative model serves as the foundation for DCM for fMRI. The first is a neural model that describes how different neuronal populations interact in a widespread network. The second links neural activity to detected hemodynamic reactions. By identifying effective connectivity between brain regions, the DCM models the fundamental dynamics of neuronal populations and how they influence one another (Friston et al., 2003).

In the state space model of DCM, we have two equations namely a neuronal equation in which we are interested and a hemodynamic equation which is the observation equation. The state space model of DCM is show in equation S1 as

$$\dot{x} = f(x, u, \theta)$$
$$y = h(x, \theta) \quad \text{(S1)}$$

Here the first equation is the neuronal equation which is the ordinary differential equation that explains the change in neuronal activity with respect to time, and second equation is the hemodynamic equation where the function h is the hemodynamic model, which specifies the biophysical processes that transform neural activity x into the Blood Oxygen-Level Dependent (BOLD) response with parameters θ.

The forward model specification is followed by the inversion of the model for each subject. The process of estimating or inversion involves identifying the parameters (like connection strengths) that strike the ideal balance between representing the data and minimizing complexity, keeping the parameters close to their previous or starting values. It is necessary to first define the priors that limit the parameters. The selected model best represents the data providing the maximu[illegible] model evidence. This is the log of the likelihood that the data set y was observed given the [illegible] m (ln p(y|m)). The negative variational free energy F is shown in equation S2.

$$\ln p(y|m) \cong F = accuracy(y, m) - complexity(m) \tag{S2}$$

spDCM is the variant of DCM. In the context of rsfMRI, spDCM uses a state space model with two components (spDCM:(Friston et al., 2014)) — a differential equation of neuronal dynamics and a hemodynamic response model — to simulate endogenous fluctuations since the external input is absent. The neuronal model of spDCM uses ‘A’ matrix only since we are focusing on resting state and there is no external stimulus. Having $x(t) = [x1(t), \dots\dots. xn(t)]^T$ is a two - dimensional array of n regions' hidden neuronal states, endogenous fluctuations, or v(t), can result from the concurrent activity of various physiological fluctuations, including those with electrophysiological, metabolic, and vasomotor origins.

$$\dot{x}(t) = A.x(t) + v(t) \tag{S3}$$

The observation error and the endogenous fluctuations represented in frequency domain as [illegible] in equation S4.

$$g_v(\omega, \theta) = \alpha_v \omega^{-\beta_v}$$
$$g_e(\omega, \theta) = \alpha_e \omega^{-\beta_e} \tag{S4}$$

Here, the complex cross spectra are represented as $\boldsymbol{g_x = X(\omega).X(\omega)^\dagger}$ where $\boldsymbol{X(\omega)}$ is the Fourier transform of $\boldsymbol{x(t)}$, $\{\boldsymbol{\alpha, \beta}\} \subset \boldsymbol{\theta}$ are the parameters governing the amplitudes and exponents of the spectral density of the neuronal fluctuations and the angular frequency is represented by $\boldsymbol{\omega = 2\pi f}$.

### Parametric empirical Bayes

Parametric empirical Bayes is a general linear model (GLM) used to measure the similarities and differences between subjects (PEB:(Friston et al., 2016; Zeidman et al., 2019)). One might want to find out whether there are any distinctions between a control group and a patient group, examine whether a drug's dosage affects certain connections, or to see whether there is a relation between connection strengths and behavioral measurements. The process requires two items from each subject θ: the covariance matrices Σ and the parameters' predicted values μ. As hemodynamics may obscure evidence for brain effects, particularly in the case of collinearity or conditional dependency, they are regarded as fixed effects and excluded from the model. PEB is a statistical model of connectivity parameters that is hierarchical. The theoretical concept of PEB can be explained with the help of equation S5 where Yi is the Observed neuroimaging data for subject, ε is the observation noise and θ are group level parameters.

$$Y_i = \Gamma_i\left(\theta_i{}^{(1)}\right) + \varepsilon_i{}^{(1)}$$

$$\theta^{(1)} = X\theta^{(2)} + \varepsilon^{(2)}$$

$$\theta^{(2)} = \eta + \varepsilon^{(3)} \tag{S5}$$

The PEB model is inverted after specifying the design matrix, and this produced two helpful values: the estimated group-level parameters and the group-level free energy. The (negative) variational free energy $F^{(2)}$, which roughly reflects the log model evidence, was also returned by inversion of the entire PEB model as a measure of its quality. It is the log of the likelihood that the neuroimaging data (from all individuals) will be observed in light of the complete hierarchical model m as shown in equation S6.

$$F^{(2)} \approx \ln p\ (Y|m) \tag{S6}$$

The free energy at the second level is equal to the DCM accuracies of all subjects added together, less the complexity brought on by fitting the DCMs and the second-level GLM. Comparing the free energy of different PEB models, the model with the greatest free energy is selected.

### Figure Legends

**Figure S1. Subject count for high, moderate, and low emotion scores**

**Figure S2. Mean and standard deviation of self-reported basic negative emotions**

**Figure S3. Accuracy of DCM model estimation**

**Figure S4. Mean connectivity matrix of self-reported high emotion scores of females (left side) and males (right side)**

(a) Mean effective connectivity matrix of self-reported high anger-affect in females (b) Mean effective connectivity matrix of self-reported high anger-affect in males (c) Mean effective connectivity matrix of self-reported high fear-affect in females (d) Mean effective connectivity matrix of self-reported high fear-affect in males (e) Mean effective connectivity matrix of self-reported high sadness in females (f) Mean effective connectivity matrix of self-reported high sadness in males. In all figures, positive value illustrates excitatory connections while negative values illustrate inhibitory connections. the diagonal values illustrate self-connections which are inhibitory.

**Figure S5. Mean connectivity matrix of self-reported low emotion scores of females and males** (a) Mean effective connectivity matrix of self-reported low anger-affect in females (b) Mean effective connectivity matrix of self-reported low anger-affect in males (c) Mean effective connectivity matrix of self-reported low fear-affect in females (d) Mean effective connectivity matrix of self-reported low fear-affect in males (e) Mean effective connectivity matrix of self-reported low sadness in females (f) Mean effective connectivity matrix of self-reported low sadness in males. In all figures, positive value illustrates excitatory connections while negative values illustrate inhibitory connections. The diagonal values illustrate self-connections which are inhibitory.

**Figure S6. Mean connectivity matrix of self-reported moderate emotion scores of females and males** (a) Mean effective connectivity matrix of self-reported moderate anger-affect in females (b) Mean effective connectivity matrix of self-reported moderate anger-affect in males (c) Mean effective connectivity matrix of self-reported moderate fear-affect in females (d) Mean effective connectivity matrix of self-reported moderate fear-affect in males (e) Mean effective connectivity matrix of self-reported moderate sadness in females (f) Mean effective connectivity matrix of self-reported moderate sadness in males. In all figures, positive value illustrates excitatory connections while negative values illustrate inhibitory connections. The diagonal values illustrate self-connections which are inhibitory.

**Figure S7. Effective connectivity associations with low anger-affect**

This figure illustrates the association between effective connectivity and low anger-affect scores in females (upper panel) and males (lower panel). In both figures the green shades represent the positive association and the red shade represents the negative association between regions. The brain plot illustrates the connections between the regions where arrowhead refers to the region that is been influenced. The solid lines show the excitatory connection whereas the dotted lines illustrate inhibitory connections. A matrix on the bottom left side shows the beta values of associations. The results are thresholded with posterior probability > 0.95. Abbreviations: mPFC = medial prefrontal cortex, PCC = posterior cingulate cortex, rHP = right hippocampus, lHP = left hippocampus, lAMG = left amygdala, rAMG = right amygdala, dACC = dorsal anterior cingulate cortex, lAI = left anterior insula, rAI = right anterior insula, lDLPFC = left dorsolateral prefrontal cortex, rDLPFC = right dorsolateral prefrontal cortex.

**Figure S8. Effective connectivity associations with moderate anger-affect**

This figure illustrates the association between effective connectivity and moderate anger-affect scores in females (upper panel) and males (lower panel). The abbreviations and color arrangement are same as in figure S7.

**Figure S9. Effective connectivity associations with low fear-affect**

This figure illustrates the association between effective connectivity and low fear-affect scores in females (upper panel) and males (lower panel). The abbreviations and color arrangement are same as in figure S7.

**Figure S10. Effective connectivity associations with moderate fear-affect**

This figure illustrates the association between effective connectivity and moderate fear-affect scores in females (upper panel) and males (lower panel). The abbreviations and color arrangement are same as in figure S7.

**Figure S11. Effective connectivity associations with low sadness**

This figure illustrates the association between effective connectivity and low sadness scores in females (upper panel) and males (lower panel). The abbreviations and color arrangement are same as in figure S7.

**Figure S12. Effective connectivity associations with moderate sadness**

This figure illustrates the association between effective connectivity and moderate sadness scores in females (upper panel) and males (lower panel). The abbreviations and color arrangement are same as in figure S7.

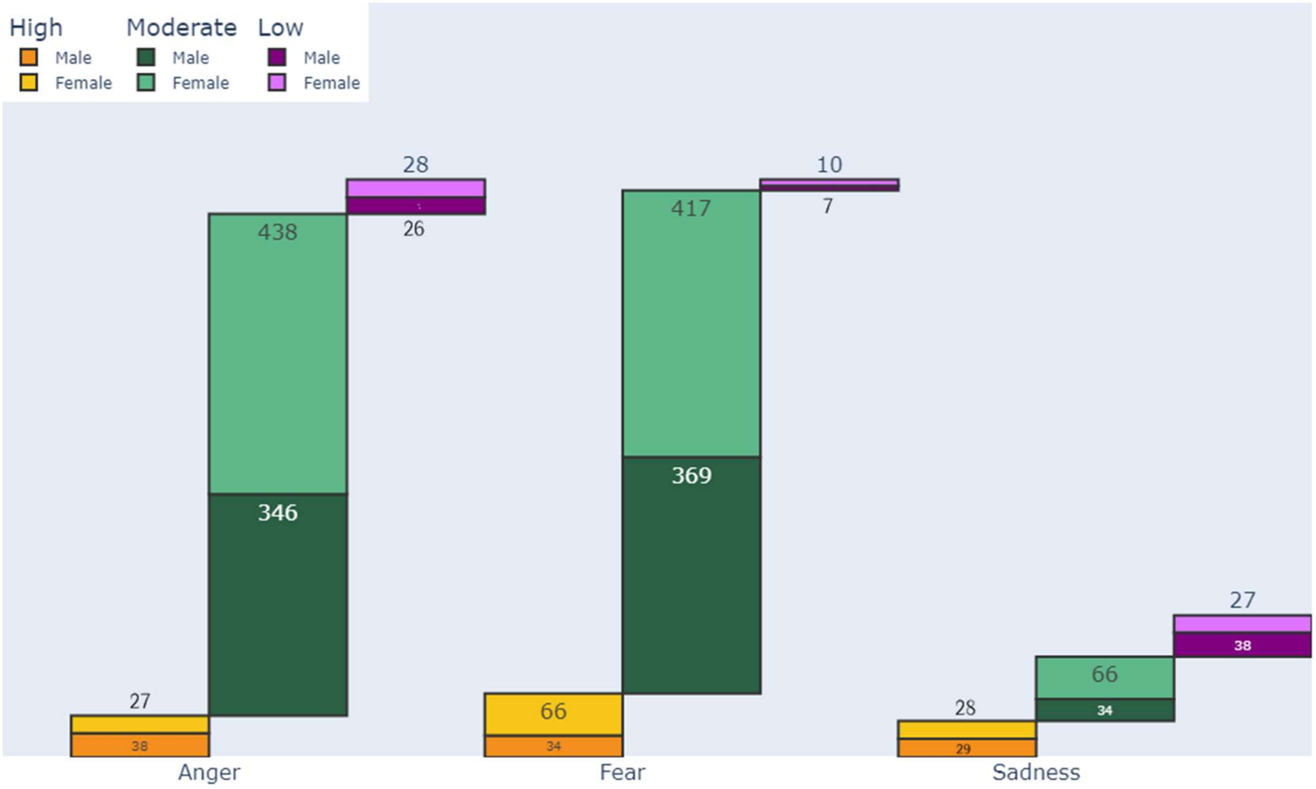


**Figure S1. Subject count for high, moderate, and low emotion scores**

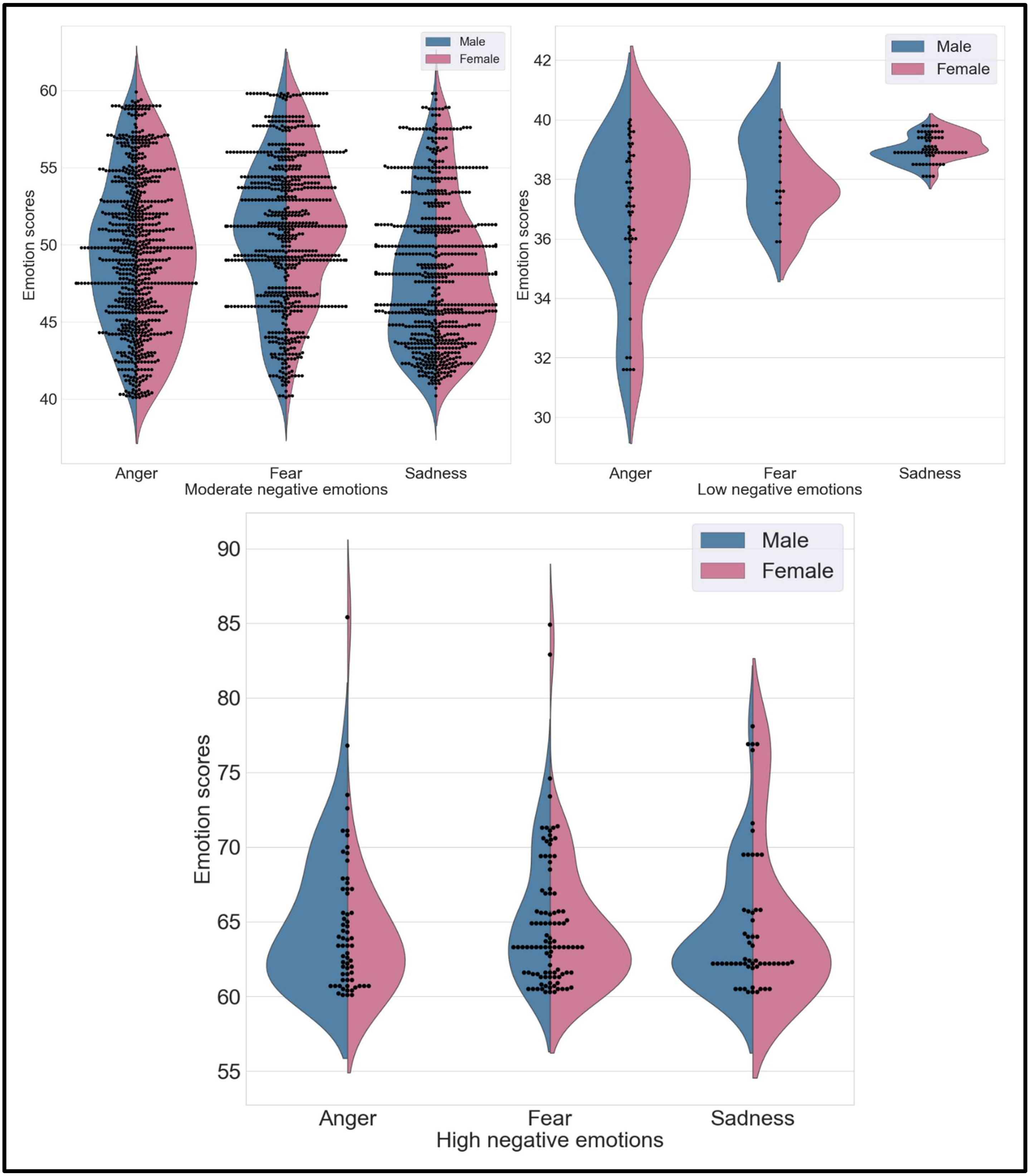


**Figure S2. Mean and standard deviation of self-reported basic negative emotions**

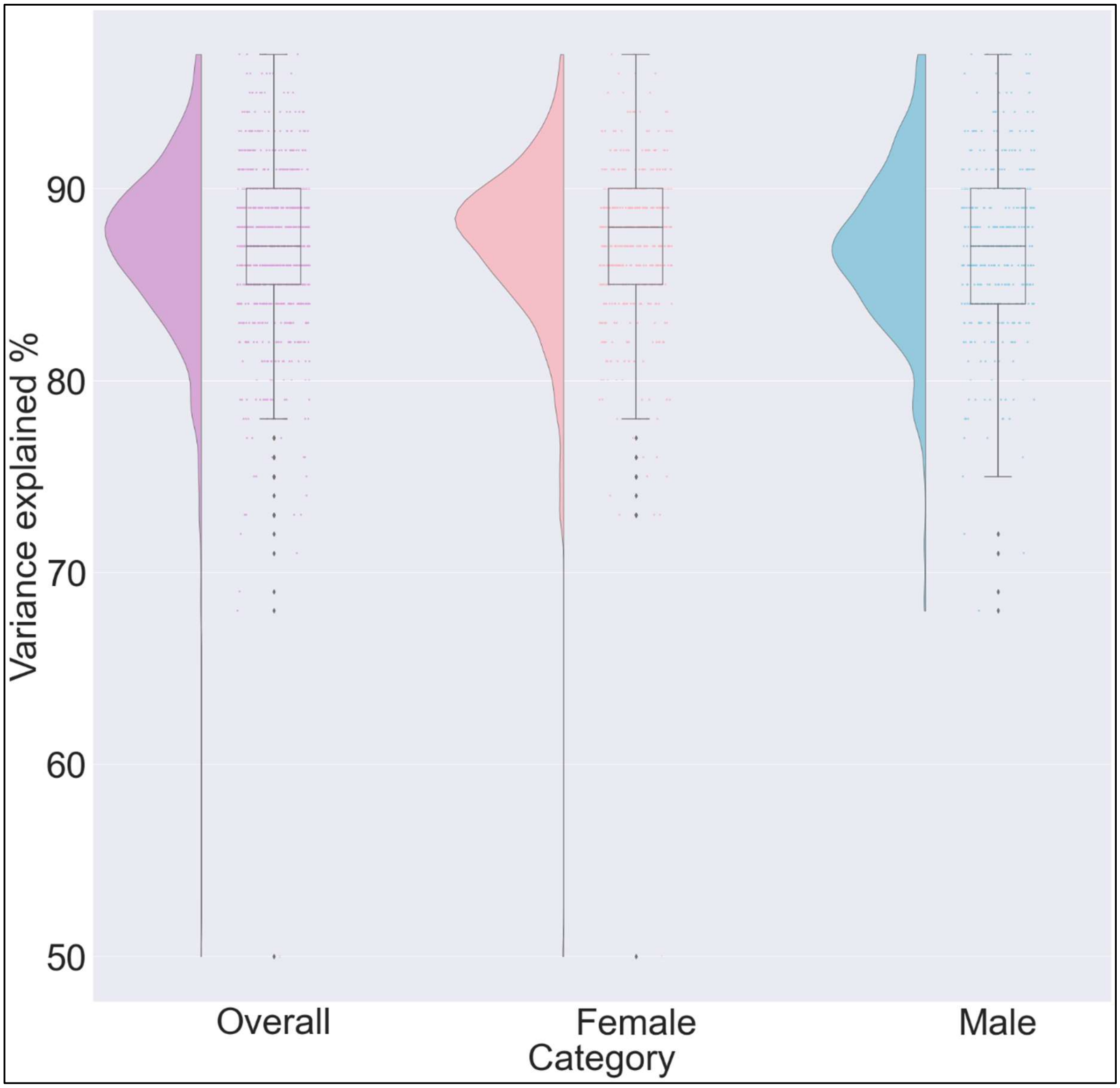


**Figure S3. Accuracy of DCM model estimation**

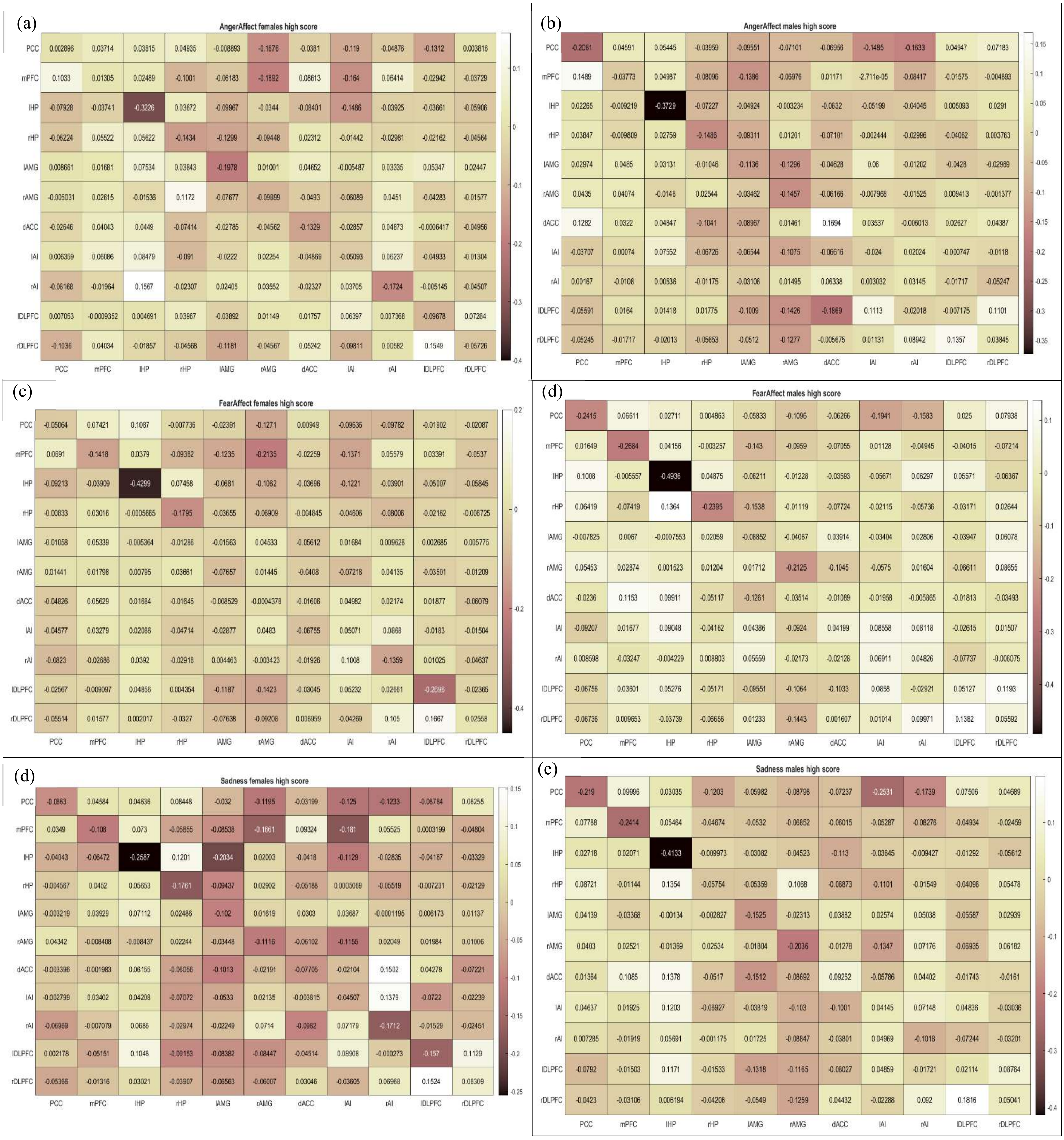


**Figure S4. Mean connectivity matrix of self-reported high negative emotion scores of females (left side) and male (right side)**

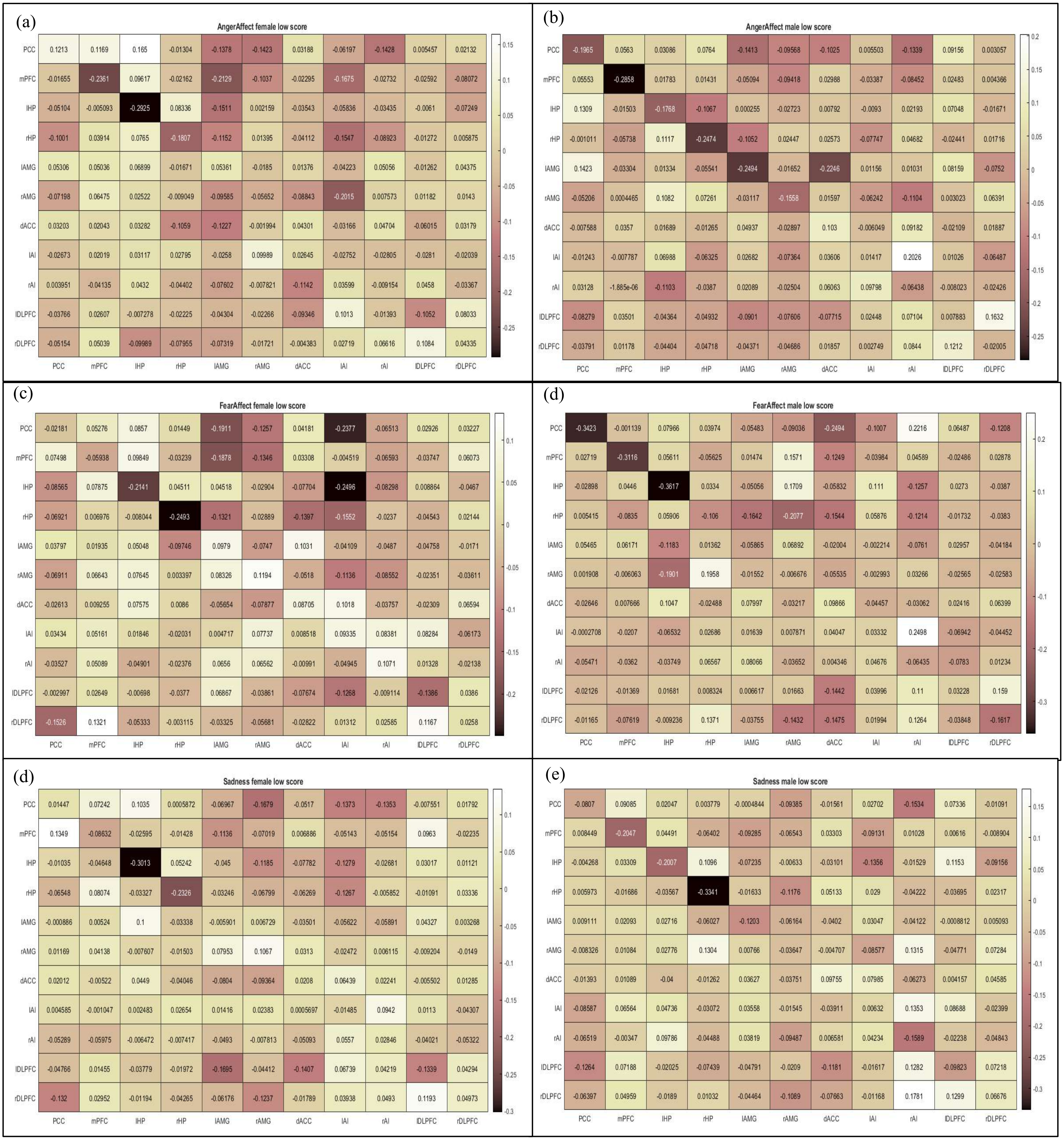


**Figure S5. Mean effective connectivity matrix of self-reported low negative emotion scores of females (left side) and males (right side)**

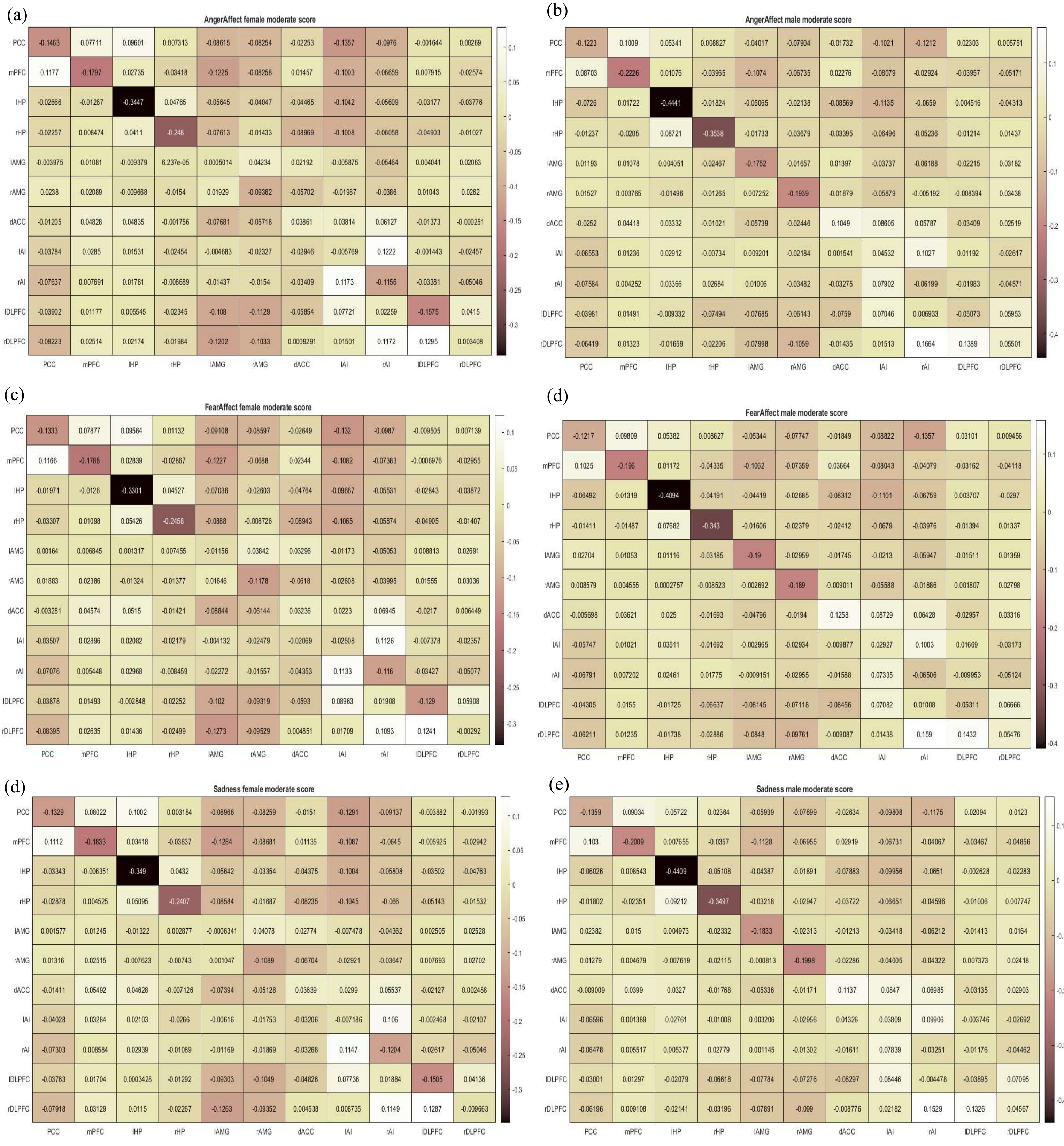


**Figure S6. Mean effective connectivity matrix of self-reported moderate negative emotion scores of females (left side) and males (right side)**

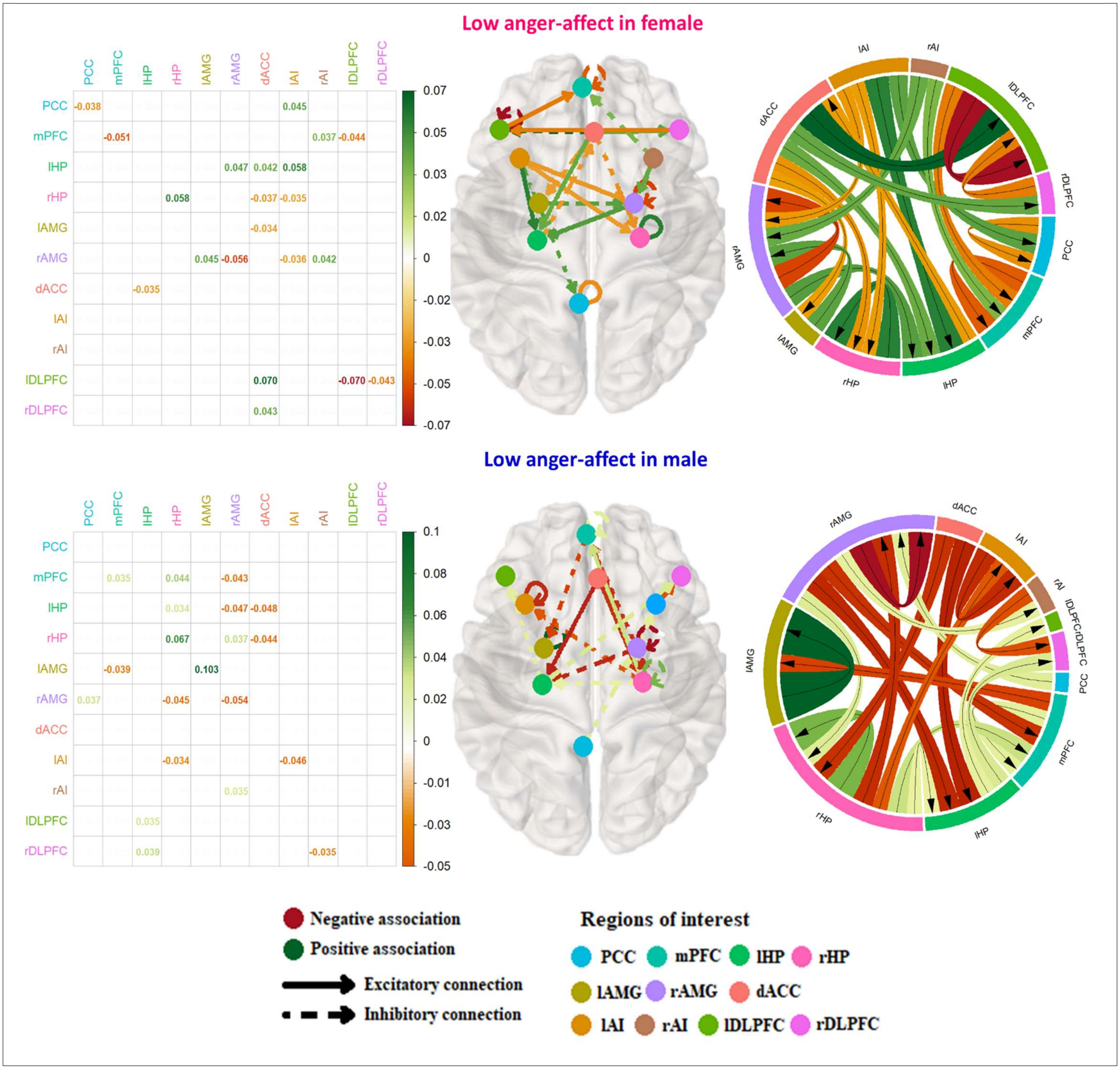


**Figure S7. Effective connectivity associations with low anger-affect**

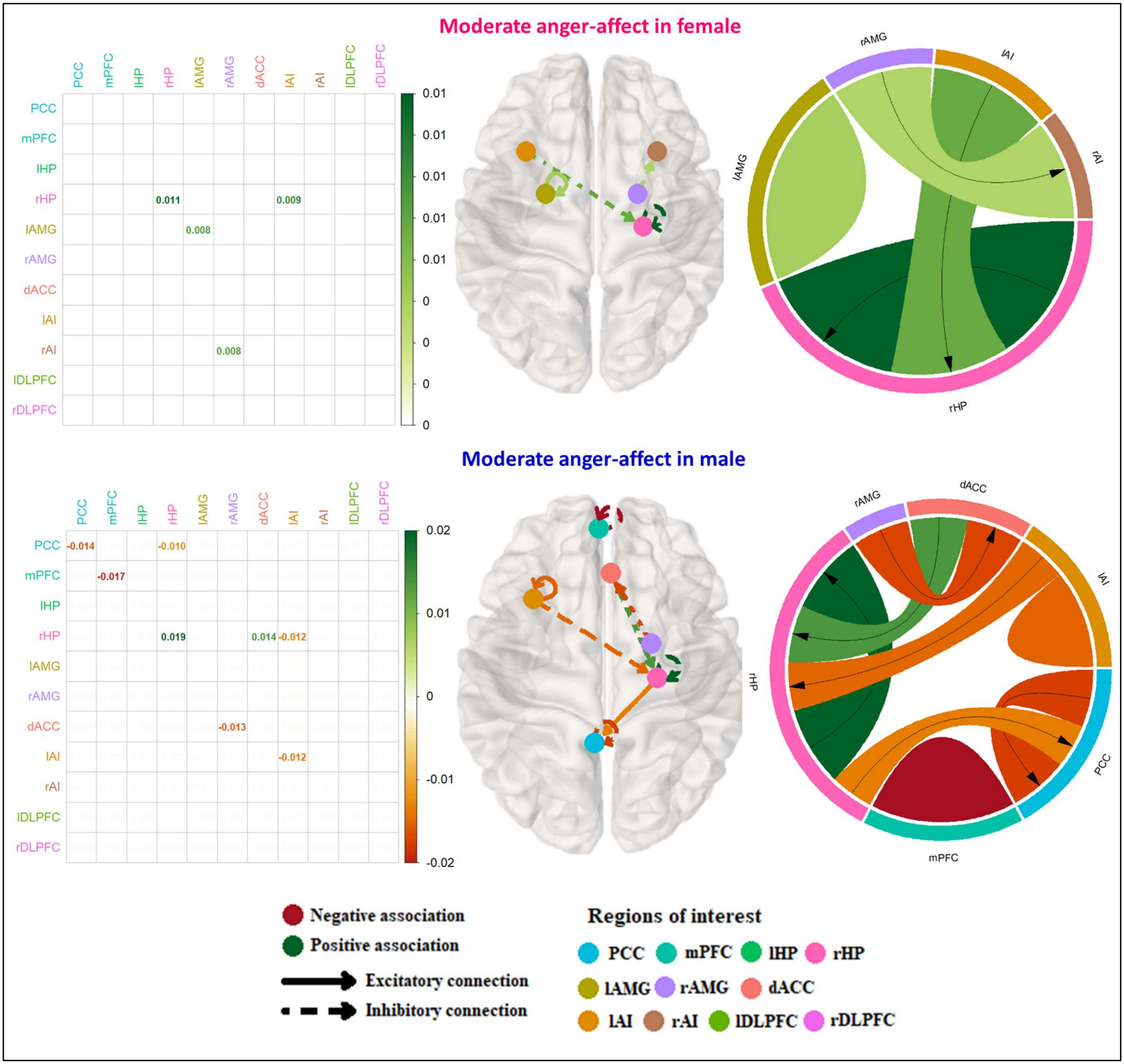


**Figure S8. Effective connectivity associations with moderate anger-affect**

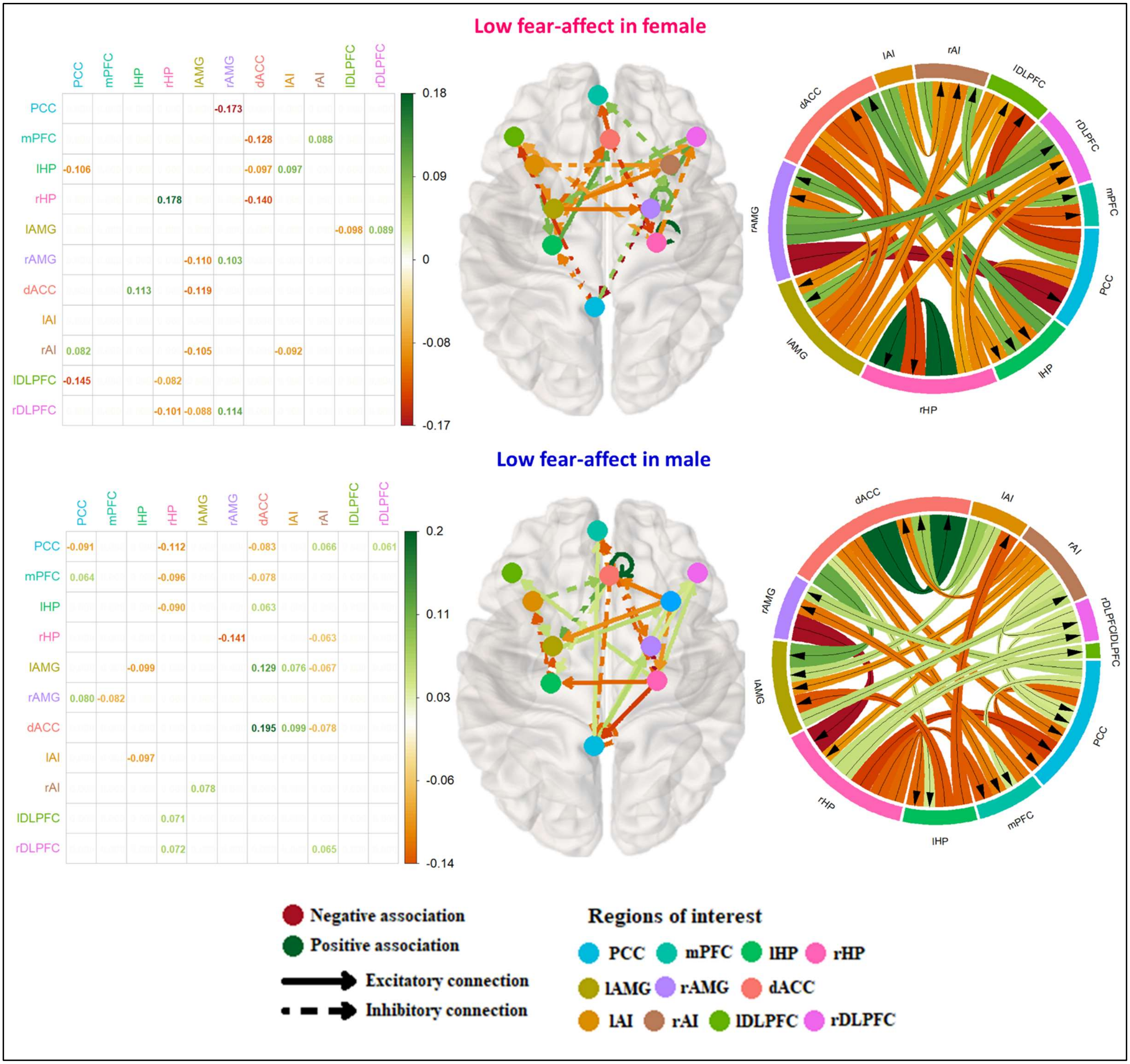


**Figure S9. Effective connectivity associations with low fear-affect**

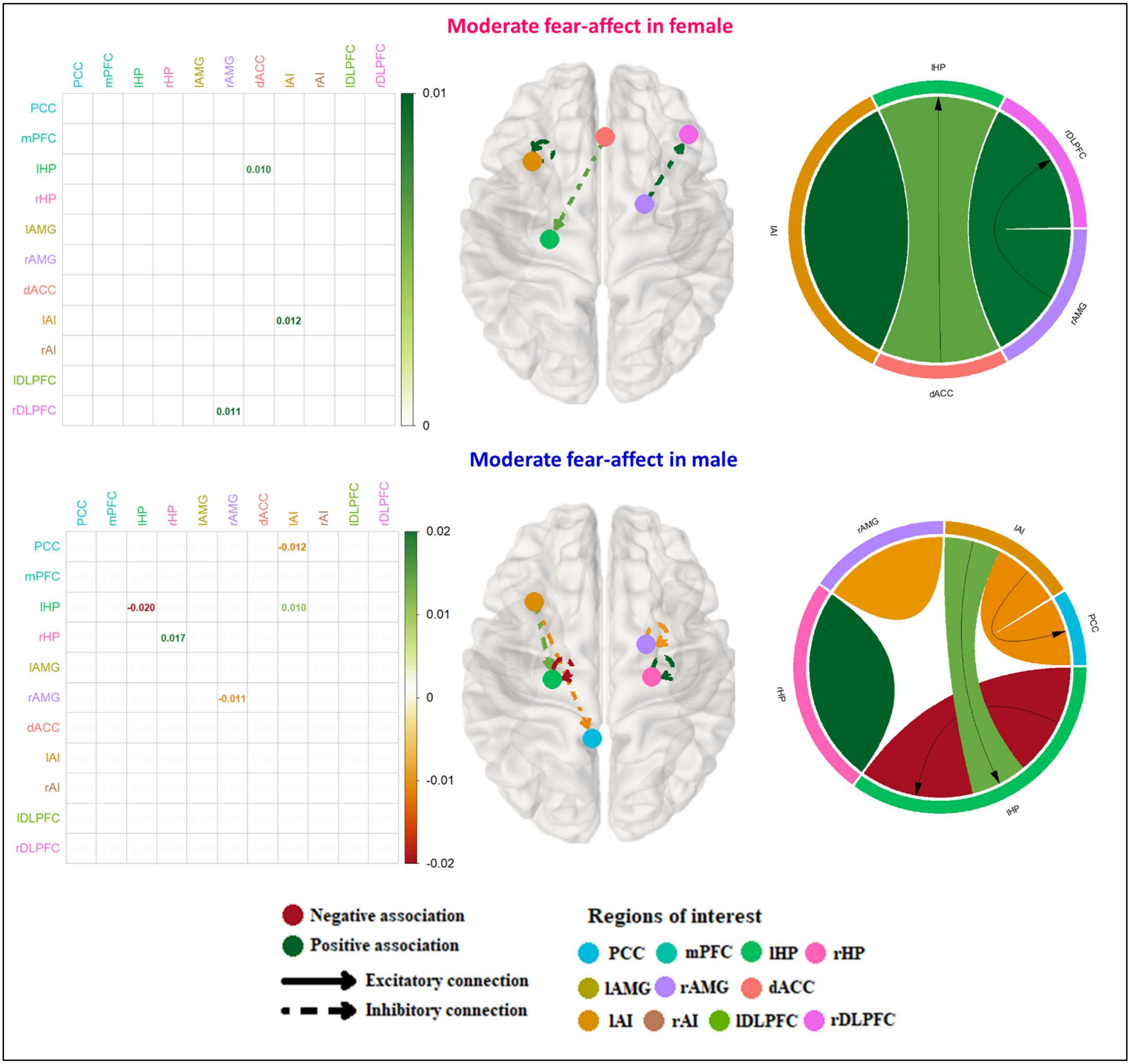


**Figure S10. Effective connectivity associations with moderate fear-affect**

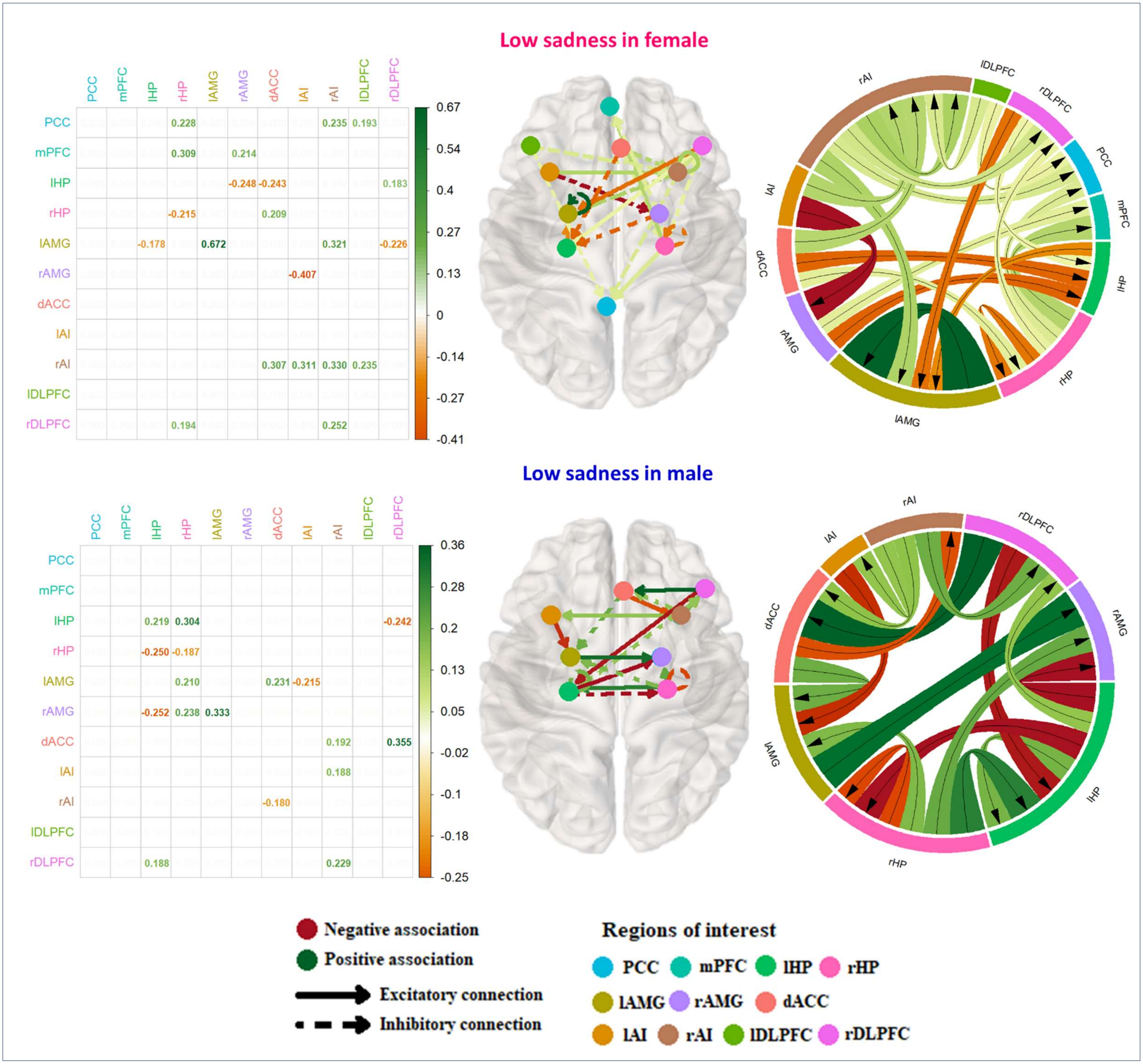


**Figure S11. Effective connectivity associations with low sadness**

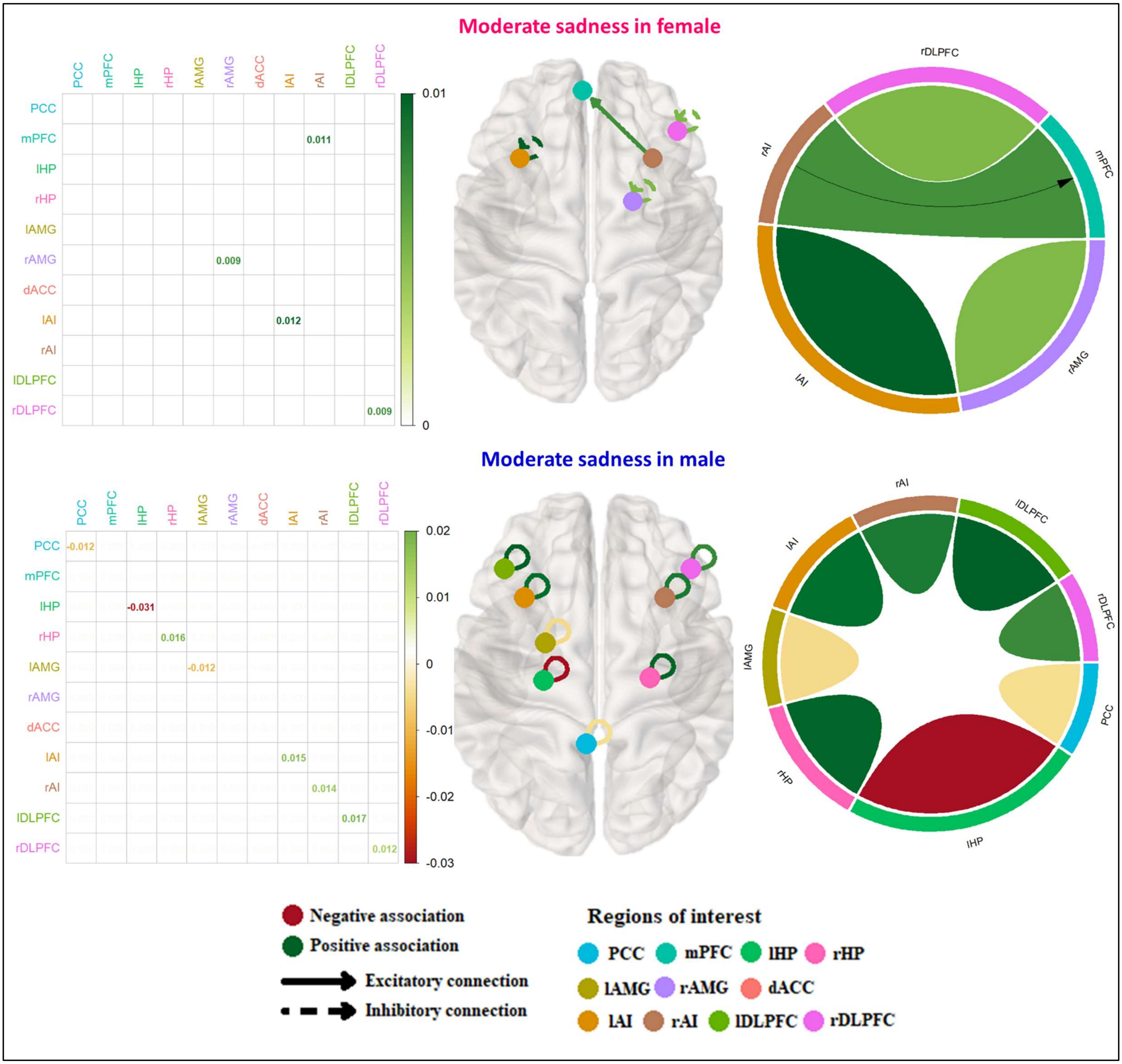


**Figure S12. Effective connectivity associations with moderate sadness**
